## Supplementary information for "Systematic decomposition of sequence determinants governing CRISPR/Cas9 specificity"

### Supplementary Figures

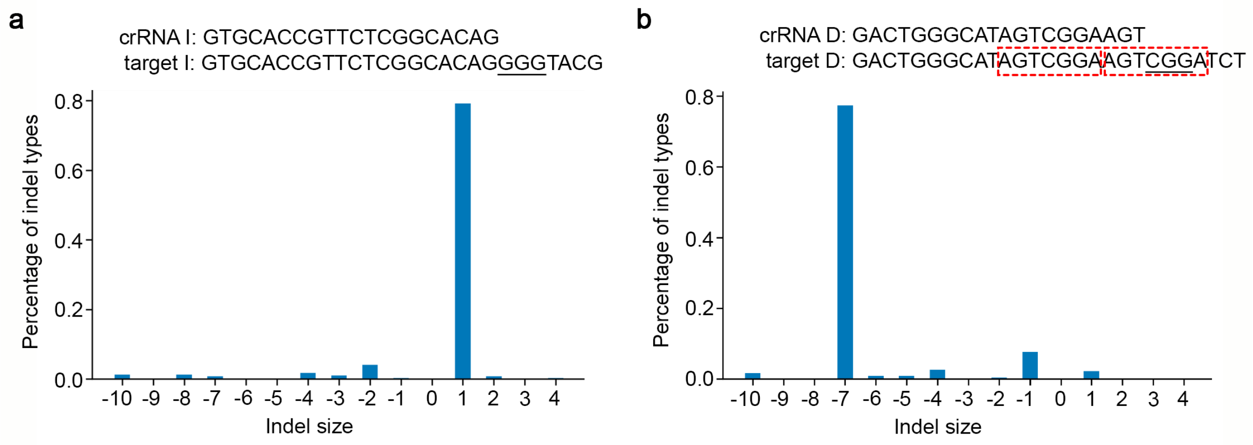


**Supplementary Figure 1**: **Indel distributions of two gRNAs for evaluation of dual-target system with distinct repair mechanisms upon double-strand breaks. (a)** gRNA I associated with dominating non-homologous end joining (NHEJ). **(b)** gRNA D associated with dominating microhomology-mediated end joining (MMEJ), where the microhomology sequences are highlighted in red rectangles. The indel distributions were calculated based on the experimental data derived from a single-target system (Allen, et al., 2018).

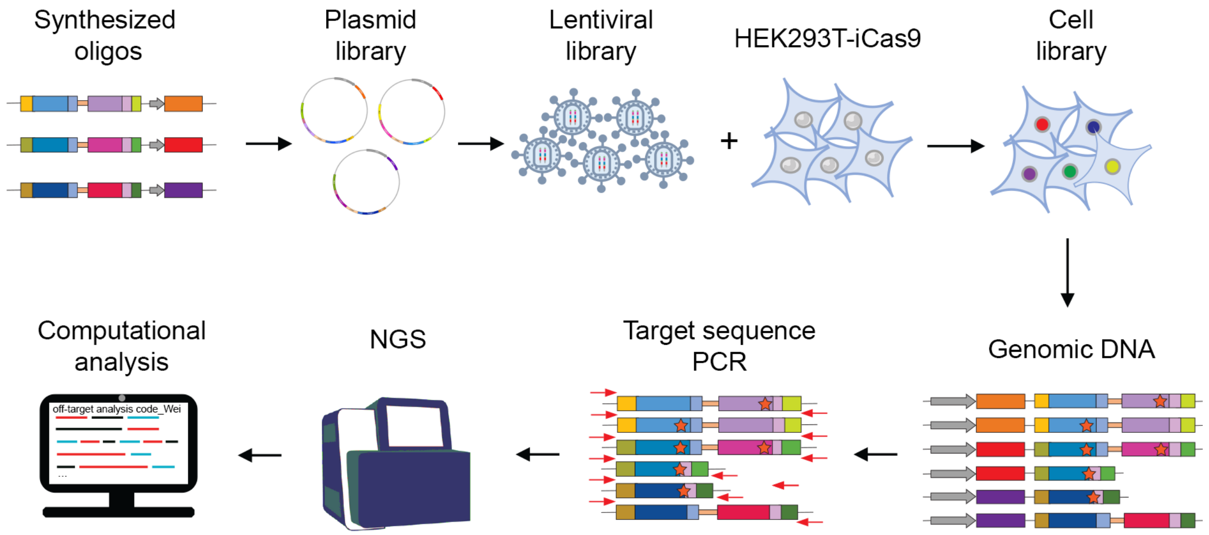


**Supplementary Figure 2**: **Schematic of experimental procedures to perform high-throughput screens.** The detailed sequence elements of each synthesized oligo library are shown in Supplementary Table 14 and Supplementary Note.

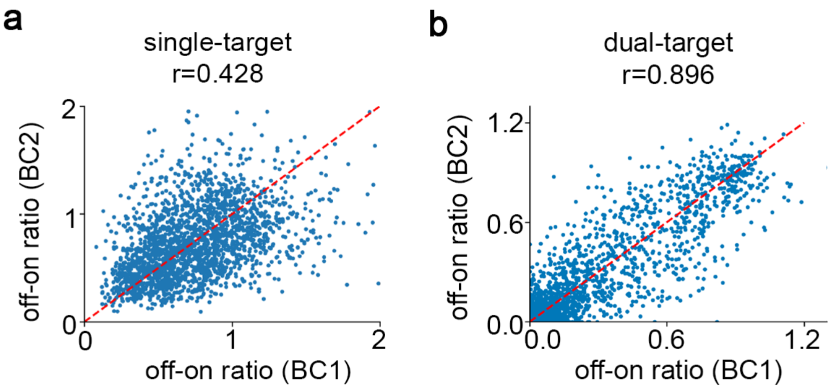


**Supplementary Figure 3**: **Correlation of off-on ratios between barcode sets.** Scatter plots showing the reproducibility of **(a)** single-target design and **(b)** dual-target design in the measurement of off-on ratios between non-overlapping barcode sets.

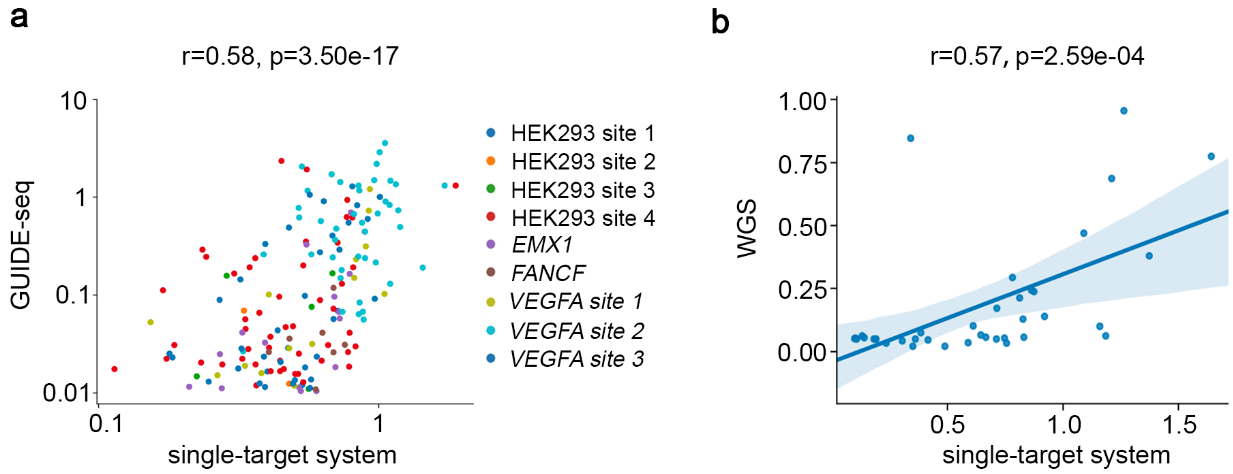


**Supplementary Figure 4: Assessment of genomic off-targets by single-target system**. Scatter plots showing the correlations between the off-on ratios estimated from the single-target system and **(a)** GUIDE-seq or **(b)** WGS, at the reported genomic off-target sequences.

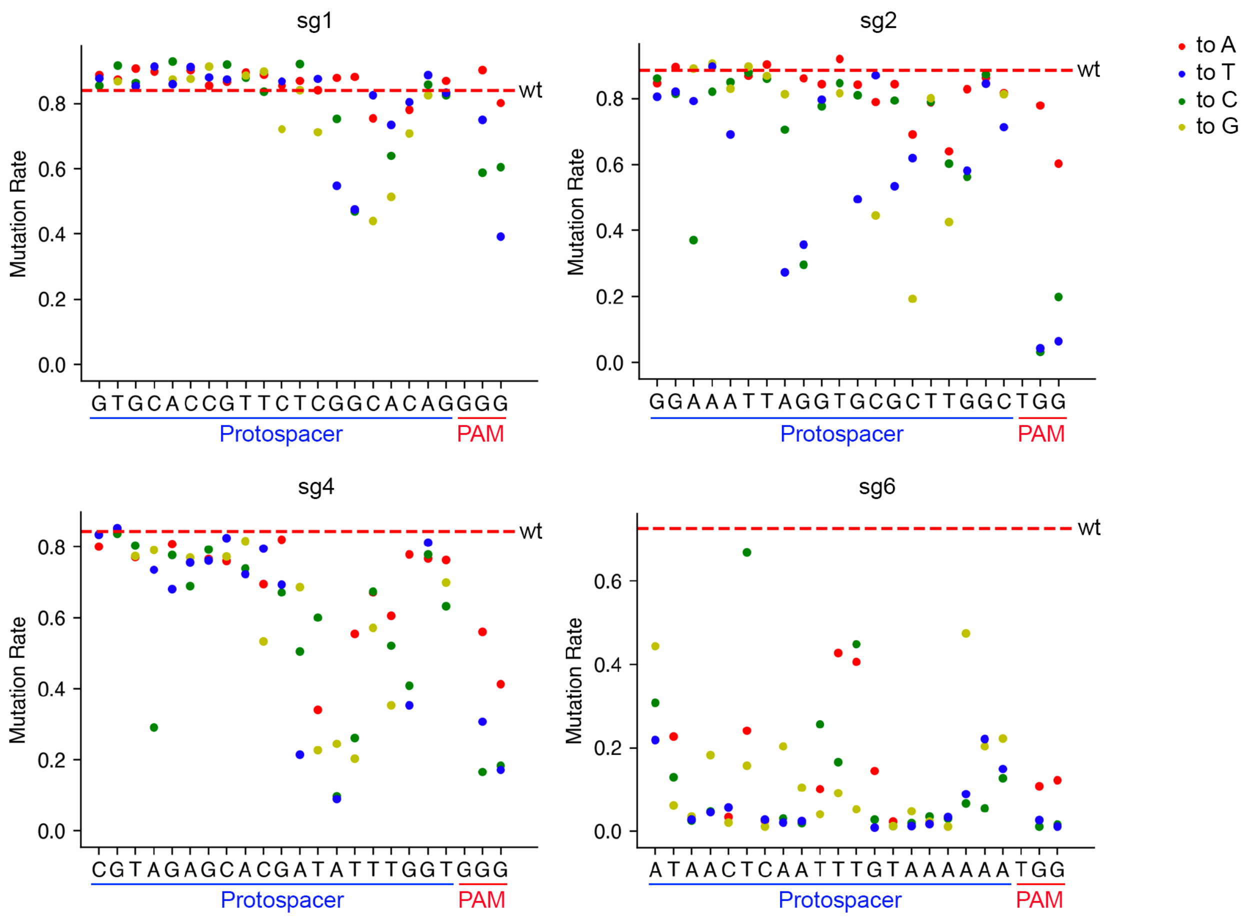


**Supplementary Figure 5**: **Single-mismatch profiles of 4 gRNAs.** The mutation rates at 1-MM target sequences of four example gRNAs that are associated with high GMT (sg1, sg2 and sg4) and low GMT (sg6). Each dot corresponds to a specific mismatched target. The red dashed line represents the mutation rate at the on-target sequence.

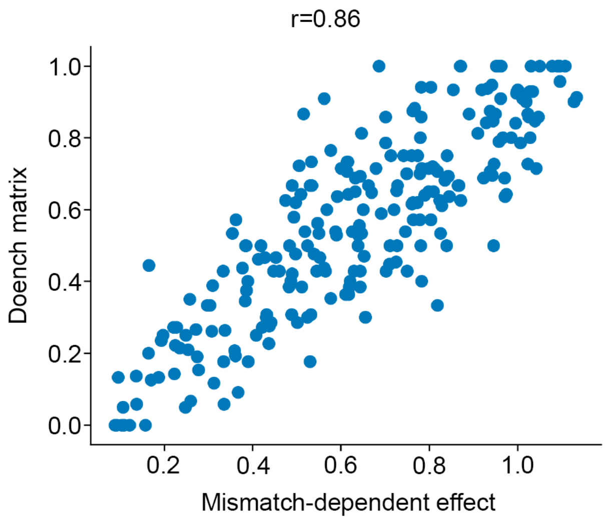


**Supplementary Figure 6**: **Correlation between 1-MM matrix and Doench's matrix**. A scatter plot showing the correlation of the mismatch-dependent effect between our and Doench's data (Doench, et al., 2016). Each dot represents the off-target effect for a certain type of single-mismatch at a specific position.

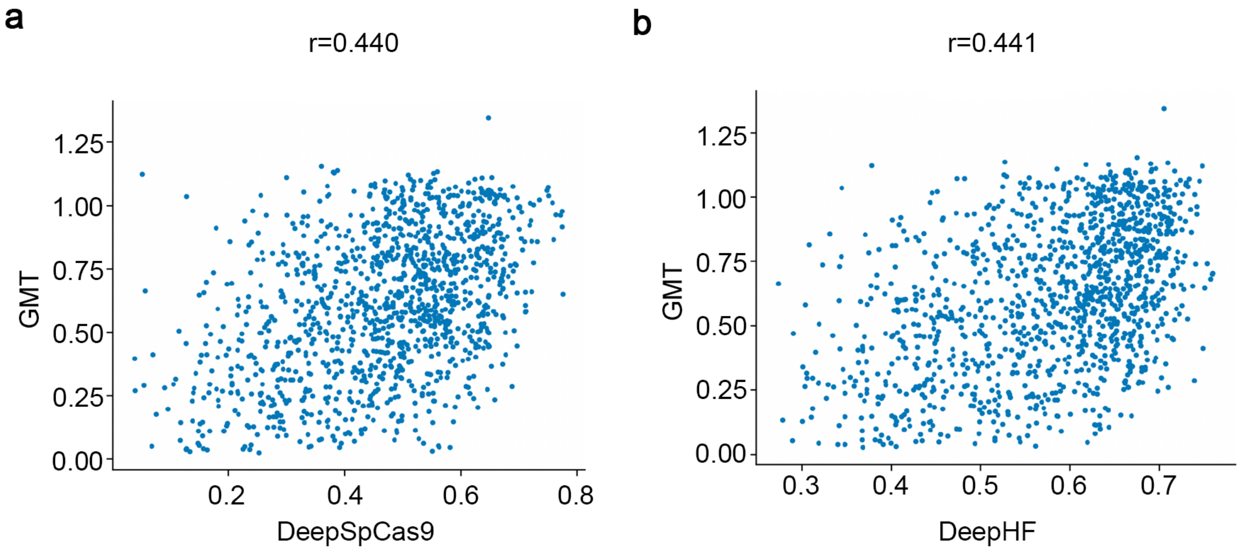


**Supplementary Figure 7**: **Correlation between GMT and gRNA activity**. Scatter plots showing the correlation between gRNA activity predicted by **(a)** DeepSpCas9 (Kim, et al., 2019) and **(b)** DeepHF (Wang, et al., 2019) and guide-intrinsic mismatch tolerance (GMT). Each dot represents a gRNA designed in the dual-target library.

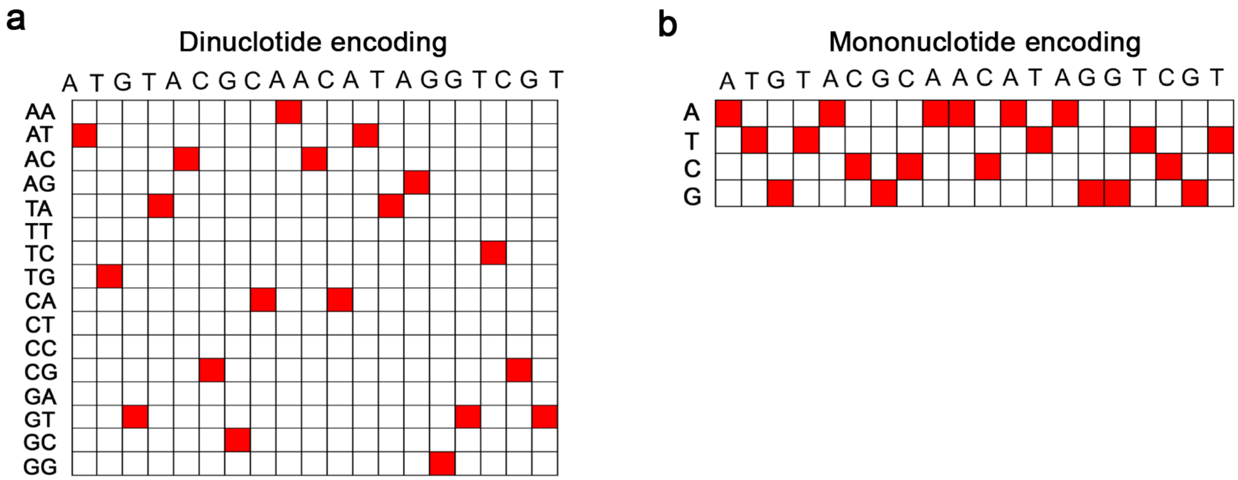


**Supplementary Figure 8**: **Schematic of two different data encoding approaches to vectorize gRNA sequence as inputs for CNN model. (a)** Dinucleotide encoding approach. **(b)** Mononucleotide encoding approach.

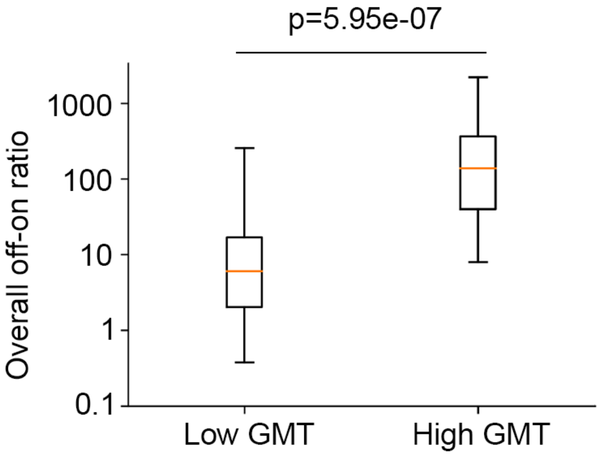


**Supplementary Figure 9**: **Comparison of the specificity between gRNAs classified into high and low GMT**. A boxplot showing the comparison of the overall off-on ratio between gRNAs with high GMT (top 25%) and low GMT (bottom 25%) in the CHANGE-seq dataset (Lazzarotto, et al., 2020), the p-value was computed using the Mann-Whitney U-test.

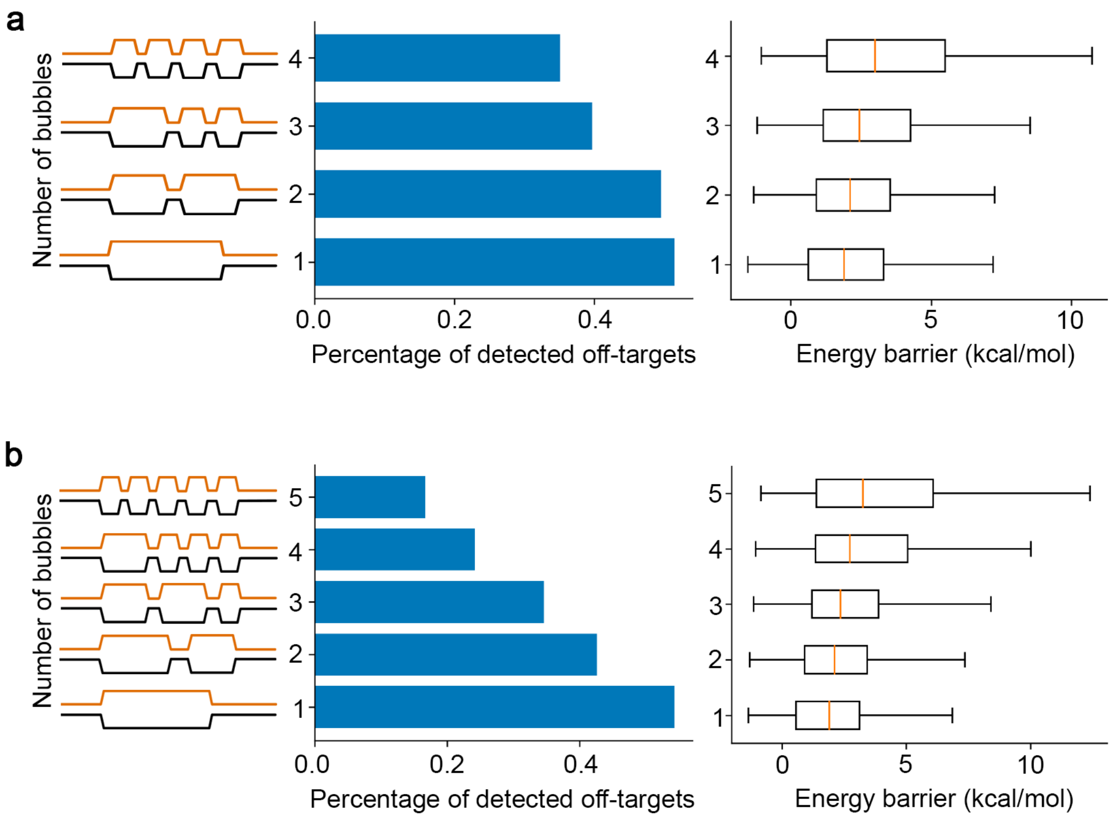


**Supplementary Figure 10**: **Correlation between bubble numbers and off-target effects.** Bar charts showing the percentage of off-target sites with different numbers of bubbles detected by CHANGE-seq (Lazzarotto, et al., 2020). Box plots showing the distribution of energy barrier computed from cumulative dinucleotide base-stacking energy changes between 1,000,000 random gRNA sequences and corresponding target sequences with randomly introduced bubbles. **(a)** Analyses on off-target sites harboring 4 mismatches. **(b)** Analyses on off-target sites harboring 5 mismatches.

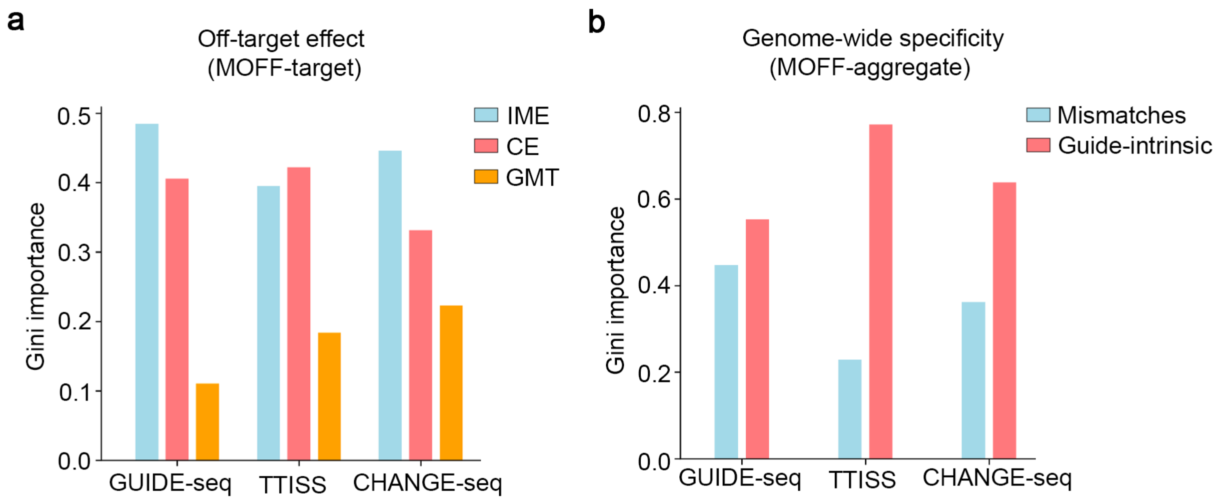


**Supplementary Figure 11**: **Comparison of feature importance in MOFF-target and MOFF-aggregate. (a)** Gini importance of IME, CE and GMT in MOFF-target. **(b)** Gini importance of mismatch-dependent effect and GMT in MOFF-aggregate.

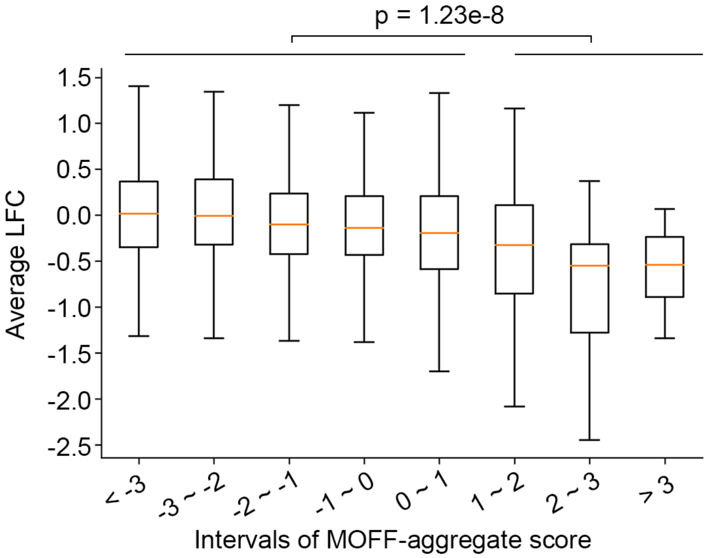


**Supplementary Figure 12**: **Correlation between MOFF-aggregate score and gRNA dropout effects in high-throughput CRISPR/Cas9 screen**. A box plot comparing the average log-fold change (LFC) of gRNAs targeting non-essential genes in GeCKO-v2 dataset across different MOFF-aggregate score intervals. The p-value was calculated using the Mann-Whitney U-test comparing the LFC of gRNAs with MOFF-aggregate scores smaller and larger than 1.

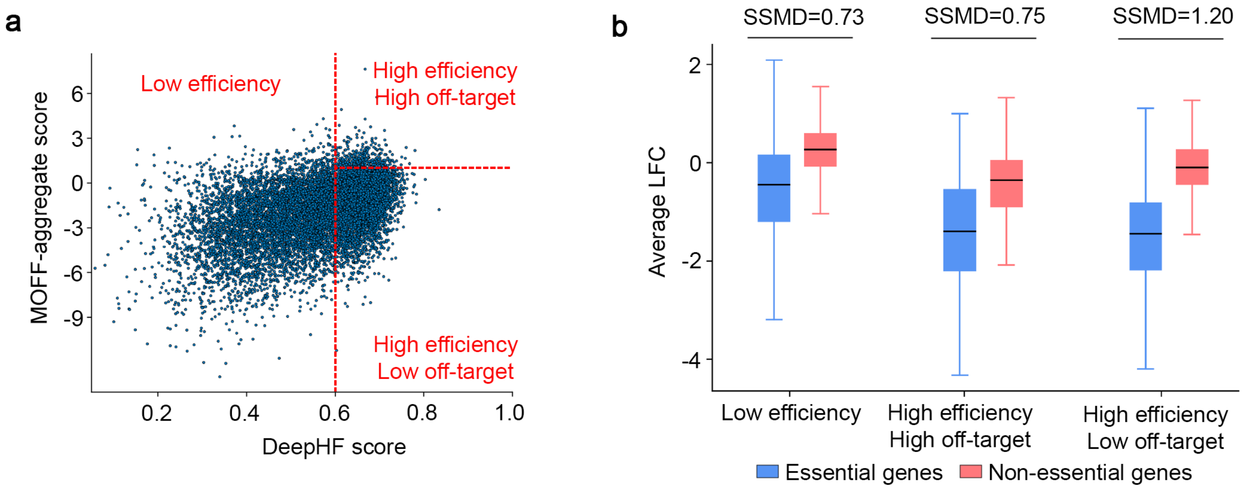


**Supplementary Figure 13**: **Application of MOFF for gRNA selection. (a)** A scatter plot showing the categorization of 11,701 gRNAs targeting 1,246 core essential genes and 758 non-essential genes in the GeCKO-v2 library, based on gRNA activity (DeepHF score, x-axis) and gRNA specificity (MOFF-aggregate score, y-axis). gRNAs were classified into 3 different categories: Low efficiency (DeepHF score < 0.6), High efficiency High off-target (DeepHF score > 0.6 and MOFF-aggregate > 1) and High efficiency Low off-target (DeepHF score > 0.6 and MOFF-aggregate < 1). **(b)** A box plot comparing the average log-fold change (LFC) of gRNAs targeting essential and non-essential genes among different gRNA categories. The differences between two groups within each category were measured by strictly standardized mean difference (SSMD).

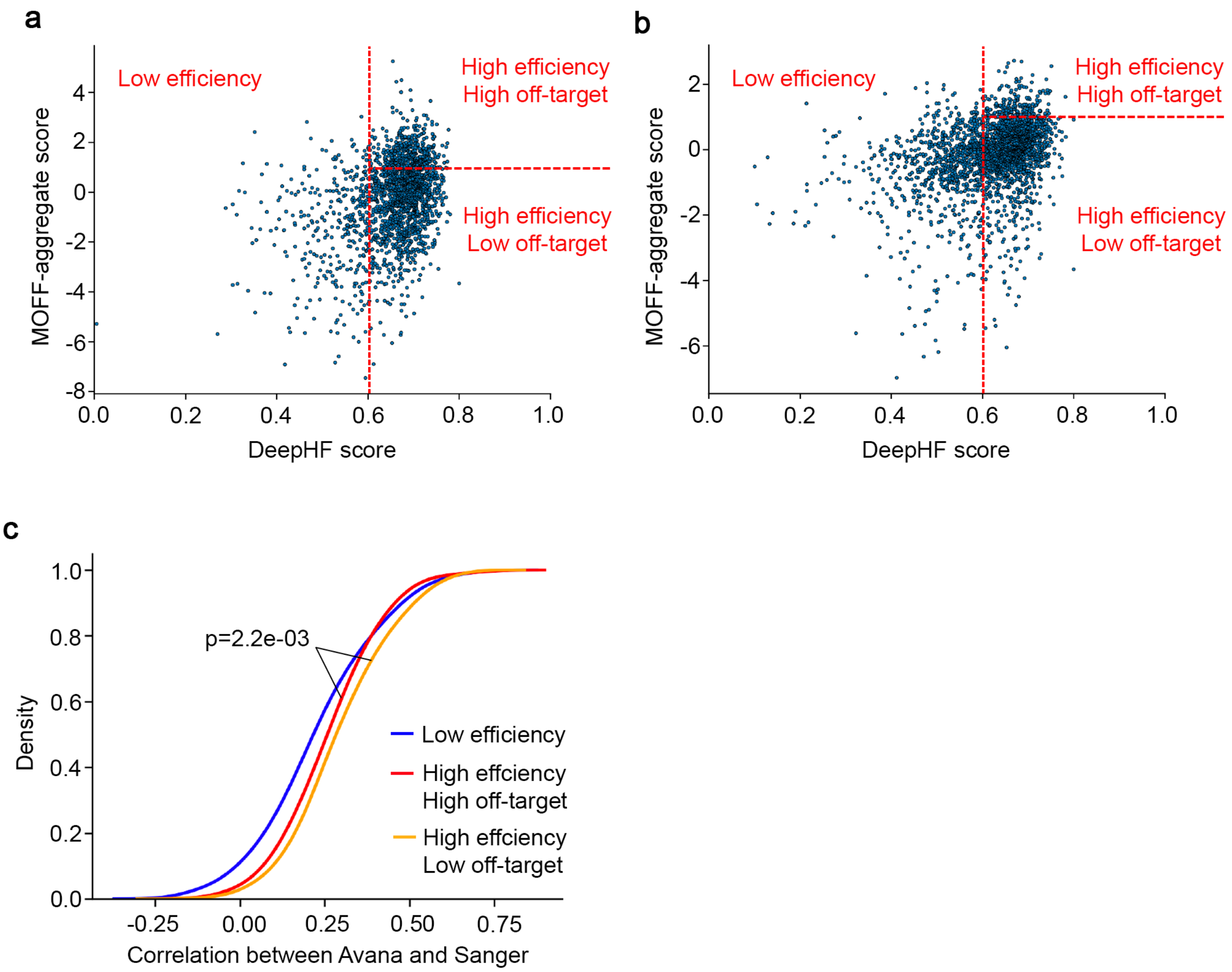


**Supplementary Figure 14**: **Comparison of the consistency between Avana and Sanger dataset among different gRNA categories. (a, b)** Scatter plots showing the categorization of gRNAs targeting 529 cell-specific essential genes in the **(a)** Avana and **(b)** Sanger libraries, based on gRNA activity (DeepHF score, x-axis) and gRNA specificity (MOFF-aggregate score, y-axis). **(c)** Cumulative distribution of correlations between gRNAs targeting cell-specific essential genes in the Avana and Sanger datasets within different categories. Pearson correlation was adopted to measure the correlation of log-fold change (LFC) caused by gRNAs from two datasets targeting the same genes across different cell lines. The p-value was calculated using the Kolmogorov-Smirnov test.

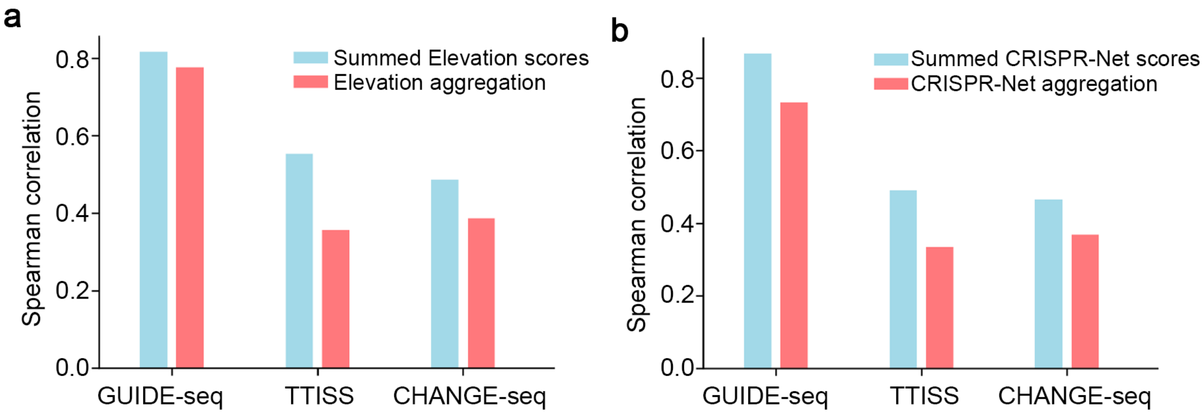


**Supplementary Figure 15: Degraded performance of machine-learning models trained on cell viability data to predict genome-wide specificity.** Bar plots comparing the performance between the sum of individual off-target score and the machine-learning based aggregation models to predict genome-wide gRNA specificity for (**a**) Elevation (Listgarten, et al., 2018) and (**b**) CRISPR-Net (Lin, et al., 2020) on three different datasets.

### Supplementary Note

**Sequence components of each oligonucleotide in different designed libraries:**

(a) Paired gRNA and dual-target libraries (T1, T2 and T3)

AAGATAGTGCAGGAACACnnnnnnnnnnNNNNNNNNNNNNNNNNNNNNNGGnnnnnnnnnnnnnnnNNNNNNNNNNNNNNNNNNNNNGGnnnnnnnnnnnnnnnAGCTTGGCGTAACTAGATCT**AATATT**GTGGAAAGGACGAAACACCgNNNNNNNNNNNNNNNNNNNNGTTTTAGAGCTAGAAATAGC

(b) Paired gRNA and single-target library (S1)

AAGATAGTGCAGGAACACnnnnnnnnnnNNNNNNNNNNNNNNNNNNNNNGGnnnnnnnnnnnnnnnnnnnnAGCTTGGCGTAACTAGATCT**AATATT**GTGGAAAGGACGAAACACCgNNNNNNNNNNNNNNNNNNNNGTTTTAGAGCTAGAAATAGC

(c) Target-only libraries (sg1, sg2, sg3, sg4, sg5 and sg6)

CTAGATCTTGAGACAAATGGTTAATnnnnnnnnnnNNNNNNNNNNNNNNNNNNNNNGGNNNNnnnnnnnnnnnnnnnnnnnnATTAACAGTATTCATCCACAATTTTAA

**Annotations for each sequence element:**

1) Primer binding sequence

2) Cloning linker sequence (containing a vector homology sequence, an SspI enzyme recognition site and an hU6 homology sequence)

3) **SspI enzyme recognition site**

4) Barcode 1

5) Barcode 2

6) Off-target sequence (Library T1, T2 and T3) / Target sequence (Library S1, sg1, sg2, sg3, sg4, sg5 and sg6)

7) On-target sequence (Library T1, T2 and T3)

8) gRNA sequence

9) Linker sequence to segregate two targets
